# Development of bright fluorescent auxin

**DOI:** 10.1101/2024.09.11.612572

**Authors:** Tsuyoshi Aoyama, Masakazu Nambo, Jia Xin Yap, Ayami Nakagawa, Marina Hayashi, Yuko Ukai, Motoo Ohtsuka, Ken-ichiro Hayashi, Yoshikatsu Sato, Yuichiro Tsuchiya

## Abstract

Polar transport of the phytohormone auxin plays a crucial role in plant growth and response to environmental stimuli. Small-molecule tools that visualize auxin distribution in intact plants enable us to understand how plants dynamically regulate auxin transport to modulate growth. In this study, we developed a new fluorescent auxin probe, BODIPY-IAA2, which effectively visualizes auxin distribution in various plant tissues. We designed this probe to be transported by auxin transporters while lacking the ability to elicit auxin signaling. Using BODIPY as the fluorophore provides bright and stable fluorescence signals, making it suitable for live-imaging under standard fluorescent microscopy. We tested the probe with auxin reporter lines in *Arabidopsis* and performed yeast two-hybrid assays. The results showed that BODIPY-IAA2 did not activate auxin signaling through the auxin receptor TIR1. However, BODIPY-IAA2 did mildly compete with both exogenous and endogenous auxins for transport, indicating that the probe is transported by auxin transporters *in vivo*. The probe not only enables visualization of its tissue distribution but also allows sub-cellular staining, including the endoplasmic reticulum and tip regions in elongating cells in moss. We also observed unusual staining patterns in the main root of non-model parasitic plants where genetic transformation is not feasible. Our new fluorescent auxin probe demonstrates significant potential for detailed studies on auxin transport and distribution across diverse plant species.

## Introduction

A phytohormone auxin regulates many plant phenomena including growth, development, and environmental responses. Auxin is polarly transported in the cell-to-cell fashion through the functions of membrane-bound auxin transporters to localize where it acts. This polar auxin transport (PAT) is dynamically changed after plants sense environmental stimuli, such as change in gravity vector or orientation of light source, to modulate their growth (Friml 2022). To understand such dynamic processes mediated by PAT, tools that visualize auxin gradient are crucial. In the past, several genetically encoded tools have been developed to visualize auxin dynamics in plants. Transcriptional reporters have been developed to visualize auxin signaling using DR5, a synthetic promoter which is transcriptionally activated by auxin (Ulmasov et al. 1997). A degron motif of AUXIN/IAA INDUCIBLE (AUX/IAA) transcription repressors, known as DII, has been utilized to report auxin-dependent degradation of fluorescent proteins through the TRANSPORT INHIBITOR RESPONSE 1 (TIR1) receptor (Brunoud et al. 2012, Liao et al. 2015). In addition, AuxSen, a Förster Resonance Energy Transfer (FRET)-type auxin sensor, is recently developed as a tool for direct auxin detection (Herud-Sikimić et al. 2021). While these reporter lines are effective in visualizing auxin distribution and its activation sites, it is difficult to analyze whether the factor is due to auxin synthesis, metabolism, or transport. Furthermore, the use of these reporter lines is limited to plants in which transformation technique is available.

In addition to these genetic tools, chemical tools have also been developed. Small-molecule phytohormones can be chemically linked to fluorescent moiety and these fluorescent hormones can be detected by fluorescent microscopy *in vivo*. These probes are available for a variety of plant hormones, including auxin, cytokinin, gibberellins, brassinosteroid, abscisic acid (Kubiasová et al. 2020, Shani et al. 2013, Irani et al. 2012, González et al. 2020). Compared with genetic tools, these fluorescent phytohormones are advantageous as they can be readily applied to any plants including those transformation is not available. In the case of auxin, fluorescein isothiocyanate (FITC) or rhodamine isothiocyanate (RITC) linked with indole-3-acetic acid (IAA), one of natural auxins, have been developed. These fluorescent auxins appeared to show both auxin-like activity and transported through PAT (Sokołowska et al. 2014). Although such probes can be expected to show similar localization to that of endogenous auxin, the effects of auxin signaling on auxin transport must be carefully considered as the expression level of auxin transporters are regulated by auxin signaling. On this idea, NBD-IAA and NBD-NAA are designed to retain functions to be transported by PAT machinery but lack the function for activating the signal transduction (Hayashi et al. 2014). These probes enabled to visualize only auxin transport activity without being affected by auxin signaling. This strategy was applied to other auxin analogs including 2,4-D which is widely used in auxin research and in tissue culture as a stable auxin analog, and also to the development of NBD-based fluorescent hormones (Pařízková et al. 2021). It is also possible to develop fluorescent auxins with anti-auxin activity (Bieleszová et al. 2019, Bieleszová et al. 2024). With the probe, phenotype caused by binding to essential proteins for auxin function can be correlated with the localization of fluorescent auxin. Both NBD and DNS-labelled IAA derivatives show anti-auxin activity at the level of PAT. While these fluorescent auxins are useful for taking still images representing auxin localization, applying it to visualize the dynamic process by live-imaging requires further improvements. For example, DNS labelled IAA derivates are unsuitable for imaging with standard fluorescence microscopy as DNS does not have suitable absorption wavelength. Although NBD is a very good fluorescent moiety due to its molecular size, it is not very bright as a fluorescent molecule. To enable live-imaging, probes with improved brightness and stability are required.

Here we developed new fluorescent auxin using BODIPY as a fluorophore. Unlike NBD-or DNS-conjugated auxins, BODIPY is bright, stable and its fluorescence spectrum is suitable for fluorescent imaging. Our results indicate that BODIPY-IAA2 has similar properties to the existing NBD-IAA, a fluorescent probe without auxin activity but retaining auxin-like transport activity. These attributes of BODIPY-IAA2 make it a useful tool for wide range of auxin studies including time-lapse observation of auxin distribution and intracellular distribution.

## Results

### Development of new fluorescent auxins

We initially synthesized a series of auxin derivatives bearing fluorescent molecules. Based on the design of NBD-IAA, alkyloxy group was selected as the linker to connect the fluorescent molecule with auxin. The alkylation of 5-hydroxy-IAA ester with bromoalkylated BODIPY units followed by hydrolysis gave the BODIPY-IAA1 and BODIPY-IAA2. Similarly, TokyoGreen, which has been reported to exhibit strong fluorescence, was also synthesized (TokyoGreen-IAA). To evaluate the specificity derived from IAA, BODIPY-indole was also prepared as a negative control.

We next tested these new fluorescent auxins by staining Arabidopsis roots. As reported previously, NBD-IAA showed fluorescent signals in root elongation zone in our microscopy set-up (Figure S1A-B). With the same treatment, BODIPY-IAA1 also showed strong signals in root elongation zone (Figure 1D). However, strong fluorescent signal was also detected in the tissue surface layer, possibly due to sticky nature of this molecules on cell wall (Figure 1D). In TokyoGreen-IAA, staining in the entire root was observed, and this probe also appeared to accumulate in vacuoles (Figure 1F and Supplemental Figure S1C). Neither of the accumulation pattern was observed in NBD-IAA. On the other hand, BODIPY-IAA2 showed similar staining pattern with BODIPY-IAA1 in the elongation zone, but staining on the surface were much more suppressed (Figure 1D). Besides, BODIPY-IAA2 strongly accumulated in ER and cytosol in the cells, which was also observed in NBD-IAA (Supplemental Figure S1A, B, D). From these results, we concluded BODIPY-IAA2 has the most similar property with NBD-IAA and decided to take this probe for further studies. To characterize physico-chemical property of BODIPY-IAA2, we first measured the spectral data of BODIPY-IAA2 (Figure 1G). Ex_max_ and Em_max_ of BODIPY-IAA2 were 496 nm and 513 nm, respectively, indicating that the fluorescent molecule has a short Stokes shift. As BODIPY is much brighter and more stable than NBD, we expected that our BODIPY-IAA2 might inherit these properties. In fact, BODIPY-IAA2 had about 60 times higher in fluorescence intensity than that of NBD-IAA (Figure 1H). These results suggest that our new fluorescent auxin BODIPY-IAA2 harbors similar characteristics to the existing NBD-IAA with improved property suitable for live-cell imaging.

**Figure 1.**
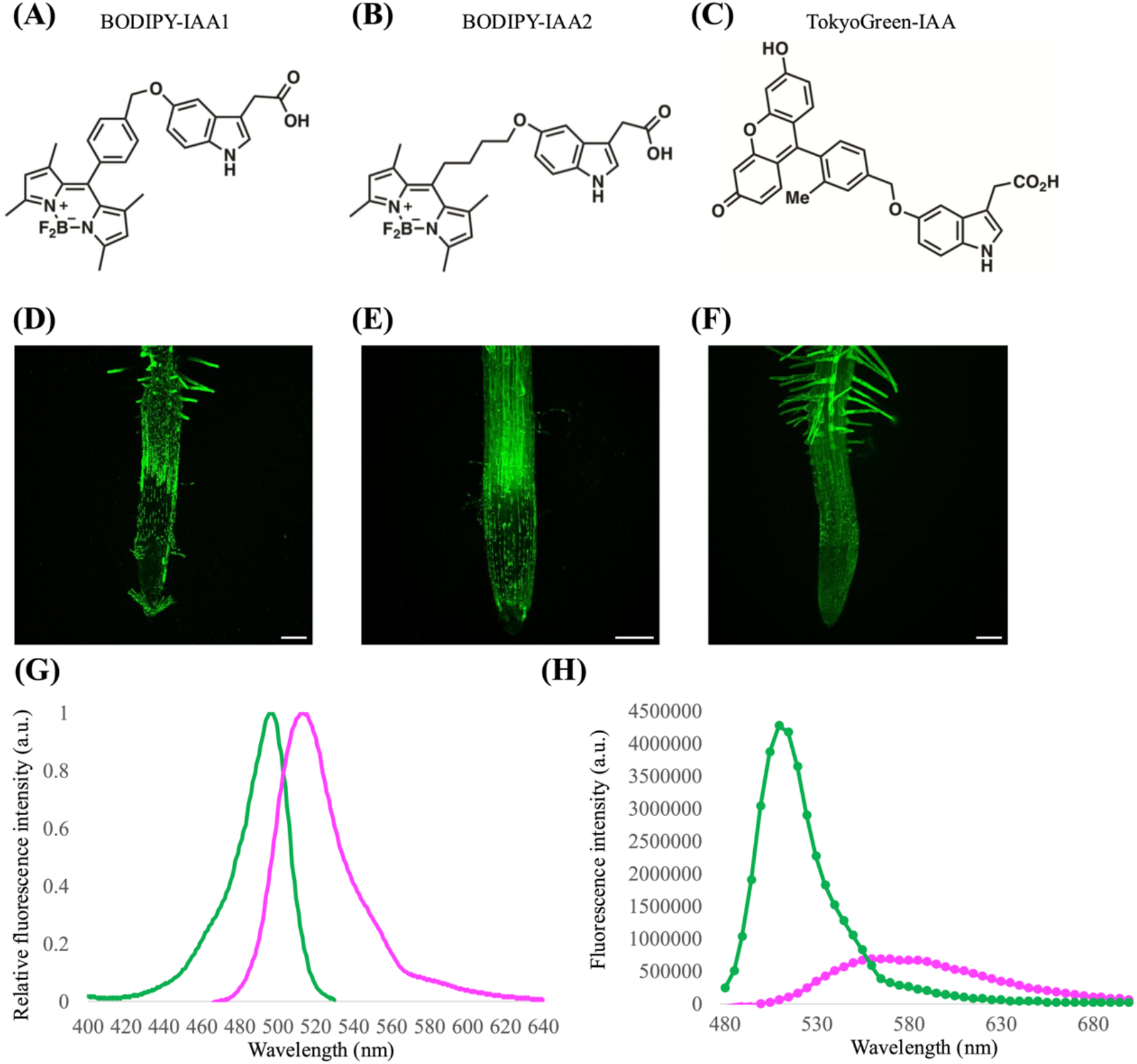
Fluorescent auxin series in this paper. **(A-C)** Structure of BODIPY-IAA1(A), BODIPY-IAA2(B), and TokyoGreen-IAA(C).**(D-F)***Arabidopsis* root was stained with 1 μM of BODIPY-IAA1(D), BODIPY-IAA2(E), and TokyoGreen-IAA(F). Scale bar: 100 μm. **(G)** Wavelength spectrum of BODIPY-IAA2. The green and magenta lines indicate the excitation and emission wavelengths, respectively. **(H)** Fluorescence intensity at various excitation wavelength in 10 μM BODIPY-IAA solution (green) and 100 μM NBD-IAA solution (magenta).

### BODIPY-IAA did not activate auxin signaling

Except for NBD-IAA, reported fluorescent auxins exhibit various auxin-like activities. As we took the design strategy of NBD-IAA and developed BODIPY-IAA2 exhibiting only auxin-like property in PAT, we next evaluated the principle using phenotype and reporter assays in *A. thaliana*. For this purpose, we took BODIPY-Indole as a negative control. In *A. thaliana*, it is well known that exogenous auxin inhibits the elongation of main roots. Consistent with this, exogenous IAA inhibited the elongation of main root in our growth condition. However, none of BODIPY-IAA2, BODIPY-Indole, nor NBD-IAA showed such an inhibitory effect on the main root (Figure 2A,B). On the other hand, BODIPY-IAA2 mildly suppressed the inhibitory effect caused by exogenous IAA, while neither BODIPY-Indole and NBD-IAA had no effect (Figure 2C). Similar results were obtained in auxin-inducible GUS reporter assays in *A. thaliana*. IAA increased the signal in the root of DR5:GUS (Ulmasov et al. 1997) and BA3:GUS (Oono et al. 1998), but none of fluorescent probes affected the staining patterns (Figure 3). On the other hand, slight suppression of GUS staining were observed in both DR5: and BA3:GUS lines when IAA was applied simultaneously with fluorescent probes (Figure 3). This result was evident in DR5:GUS, in which dose-dependent reduction of GUS staining was observed in BODIPY-IAA2 (Supplemental Figure S2, S3). These results indicate that BODIPY-IAA2 doesn’t activate auxin signaling, but competes with IAA for uptake into roots. We further confirmed the results by yeast two-hybrid (Y2H) assay. The auxin receptor, TIR1, interacts with its downstream signaling component called IAA proteins after binding to auxin, and this auxin-dependent protein-protein interaction can be detected as blue staining representing the expression of *LacZ* reporter (Uchida et al., 2018). While we observed blue staining in response to IAA, none of fluorescent probes induced Y2H interaction at 100 µM (Figure 4). Up to 1 µM, BODIPY-IAA2 didn’t compete with 0.1 µM IAA in this Y2H assay, but slight reduction of blue staining was observed at 10 µM. These Y2H results suggests that BODIPY-IAA2 doesn’t activate auxin signaling, but compete with IAA uptake in yeast too. In summary our results suggest that BODIPY-IAA2 has no activity on activating auxin signaling, but possibly competes with auxin for transports.

**Figure 2.**
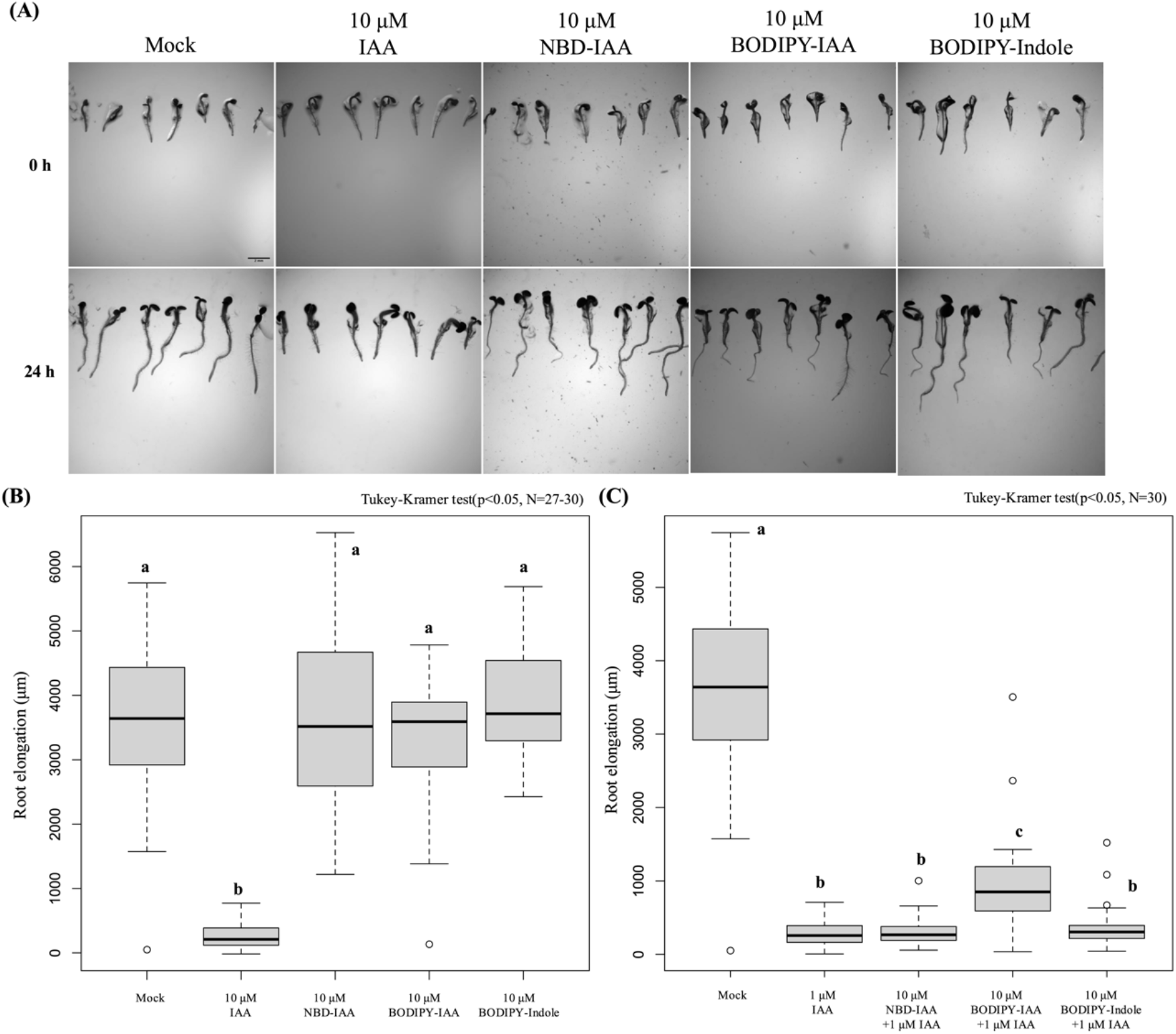
Root elongation assay using fluorescent probes. **(A)** Three-day-old *Arabidopsis* seedling grown on fluorescent auxins. Root length was measured after 0h and 24h incubation on 1/2 MS agar medium containing 10 µM auxin or fluorescent auxins. Scale bar: 2 mm. **(B, C)** Root length of *Arabidopsis* seedlings in (A). The different alphabets indicate that the results were statistically significant.

**Figure 3.**
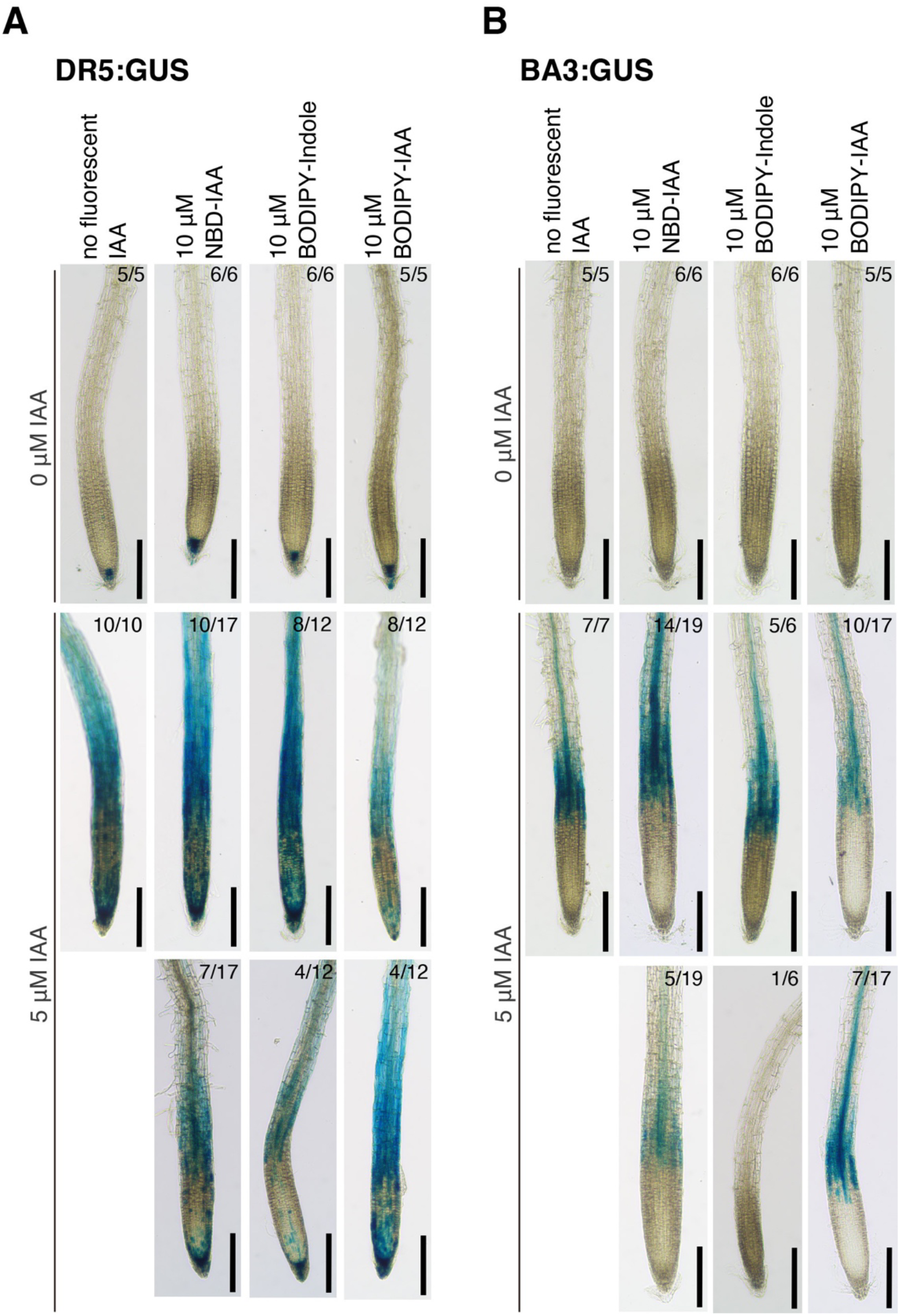
Effects of Fluorescent probes on auxin-responsive reporter gene expression. Six-day-old *Arabidopsis* DR5:GUS (A) and BA3:GUS (B) seedlings were incubated on 0 or 5 μM IAA with NBD-IAA, BODIPY-Indole or BODIPY-IAA for 4 h. Bar = 0.2 mm. Fractions on each image indicate the number of phenotypes observed (numerator) and the number of seedlings examined (denominator).

**Figure 4.**
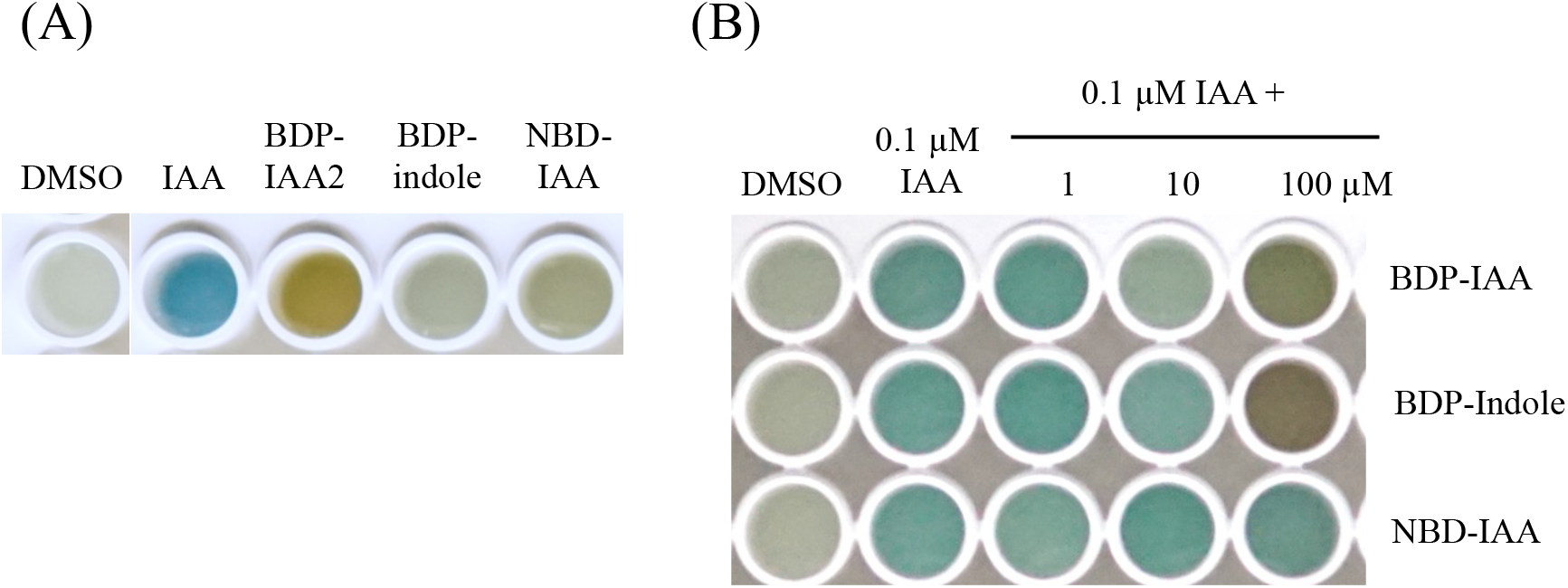
Effects of Fluorescent IAAs on Y2H interaction between TIR1 and IAA7. **(A)** X-gal staining indicating yeast two hybrid interaction between auxin receptor TIR1 and IAA7 in response to 100 µM IAA or fluorescent auxins. **(B)** Competition of fluorescent probes with 0.1 µM IAA.

### Distribution of fluorescent auxin BODIPY-IAA in *planta*

We next evaluated whether BODIPY-IAA2 visualizes *in vivo* auxin distribution in various plants and tissues. Firstly, we observed BODIPY-IAA2 localization in the main root of *Arabidopsis* where distribution of auxin is well-studied. As shown in Figure 5A, BODIPY-IAA2 showed strong signals in the root cap, elongation zone, and root hair. This staining pattern is similar to that observed with NBD-IAA (Figure 5C, Hayashi et al., 2014). In addition, we were unable to detect BODIPY-Indole signal deep in root tissue, while the BODIPY-IAA and NBD-IAA staining was detected in internal tissue of the root (Figure 5D-F, Supplemental movie1-3). These data suggest that localization of BODIPY-IAA2 relies on the IAA portion of the molecule rather than the BODIPY. We noticed that staining patterns of BODIPY-IAA2 in the lateral roots were different from that in the main root, as strong signal was detected around the root tip throughout whole developmental processes (Figure 5G-I). Overall, the pattern of BODIPY-IAA2 localization is consistent with NBD-IAA and previously reported auxin distribution in *A. thaliana*.

**Figure 5.**
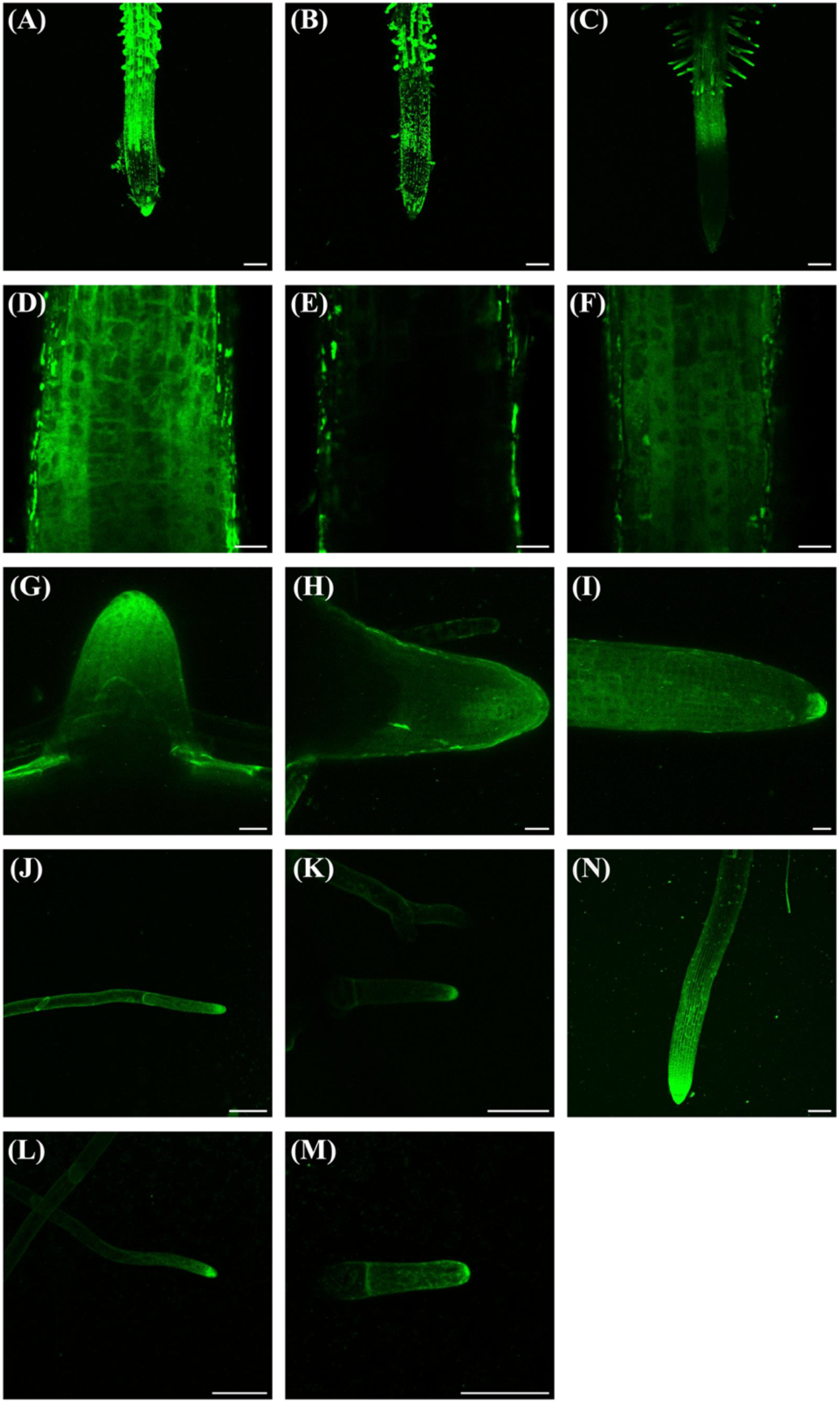
BODIPY-IAA2 staining using various plants. **(A-N)** Z-stack images taken by confocal microscopy. Maximum intensity projection images were shown (A-C, G-N). **(A-F)** Main roots of *Arabidopsis* were stained with 1 μM BODIPY-IAA2 (A, D), BODIPY-Indole (B, E), and NBD-IAA(C, F). Scale bar: 100 μm (A-C), 20 μm (D-F). **(G-I)** Lateral roots of *Arabidopsis* stained with 1 μM BODIPY-IAA2. Scale bar: 100 μm. **(J-M)** Caulonema (J,L) and chloronema (K,M) cells of *P. patens* stained with 1 μM BODIPY-IAA2 (J, K) and NBD-IAA (L, M). Scale bar: 50 μm. **(N)** Roots of *P. viscosa* stained with 1 μM BODIPY-IAA2. Scale bar: 100 μm.

To explore possible utility of this probe in different plant species, we next performed staining the moss *Physcomitrium patens*. We detected strong BODIPY-IAA2 signal in apical cells in both caulonema and chloronema cells (Figure 5J-M). Interestingly, BODIPY-IAA2 localized strongly to the tip region in apical cells (Figure 5J, K). This intracellular gradient of fluorescent auxin could be related to the function of auxin in protonema cells of the moss, and our new probe could be used to track such intracellular auxin distribution and function. Similar patterns of staining were obtained from NBD-IAA, but BODIPY-IAA2 showed much brighter images. Finally, we applied our BODIPY-IAA to non-model parasitic plants called *Parentucellia viscosa*. Unlike genetically encoded fluorescent reporters, our BODIPY-IAA2 can be used in non-model plants, such as *P. viscosa*, in which transformation technique is not available. Interestingly, the staining pattern of the main root in *P. viscosa* was similar to that of the lateral roots in *Arabidopsis* as strong signal was observed at the root tip (Figure 5N). These results suggest that BODIPY-IAA2 can be used to detect differential regulation of PAT among plant species.

### Characteristics of BODIPY-IAA as fluorescent auxin

If BODIPY-IAA2 is transported through PAT machinery, high concentration of the molecule might compete with endogenous auxin for auxin transporters. To test the idea, we performed root gravitropic response assay as it is well studied that PAT is required for root gravitropism (Figure 6). As expected, we observed no effect of BODIPY-IAA2 at low concentration (2 µM) corresponding to our optimal treatment for staining (Figure 6C, I), while root response to gravity was lost at 50 µM in a similar way to auxin transport inhibitor NPA (Figure 6B, F, J). The similar effect was also observed in high concentration of NBD-IAA (Figure 6E, H, I, J). On the other hand, BODIPY-Indole did not affect gravitropism in either condition, suggesting again that the BODIPY itself wasn’t responsible for the effect (Figure 6D, G, I, J). These results support the idea that BODIPY-IAA2 is transported by auxin transporters and competes with endogenous auxin during root gravitropism.

**Figure 6.**
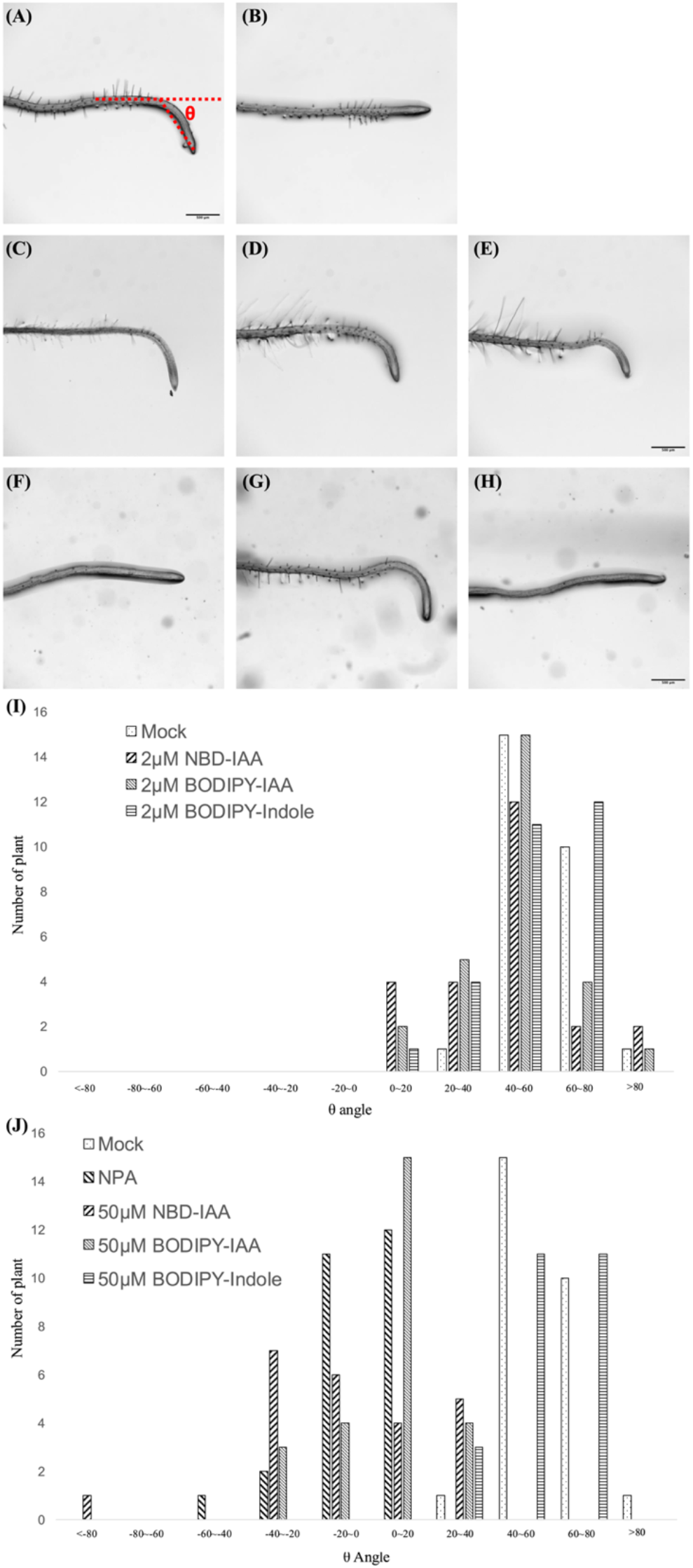
Root gravitropism assay. **(A-H)** Five-day-old *Arabidopsis* seedling incubated for 6h at 90 degree tilt in dark condition with each chemicals: Mock (A), 2 μM NPA (B), 2 μM BODIPY-IAA (C), 2 μM BODIPY-Indole (D), 2 μM NBD-IAA (E), 50 μM BODIPY-IAA (F), 50 μM BODIPY-Indole (G), and 50 μM NBD-IAA (H). Scale bar: 500 μm. **(I-J)** The degree to which the roots exhibited the bending during 6 h of incubation was quantified as θ (θ in (A)).

To further confirm the idea, we tested the effect of auxin transport inhibitors on the staining pattern of BODIPY-IAA2 (Figure 7). 1-NOA is an inhibitor that targets auxin influx carrier AUXIN RESISTANT 1/LIKE AUXIN RESISTANT (AUX1/LAX). On this inhibitor, staining of BODIPY-IAA2 was strong and uniform in the lateral root cap and epidermis (Figure 7B, F, Supplemental movie5). On the other hand, the signal was weak in the rest of cells. These results suggest that uptake of BODIPY-IAA2 from media is independent from these auxin importers, but transport into inner tissues requires AUX1/LAX. Similar results were observed with TIBA, a widely used auxin transport inhibitor with no clear mechanism of action revealed (Figure 7C, G, Supplemental movie6). The strong uniform signals in the lateral root cap and epidermal cells in the presence of TIBA suggest that auxin transporters are not involved in the uptake of BODIPY-IAA2 into these tissues. Bz-NAA is an auxin transport inhibitor which targets auxin efflux carrier PIN-FORMED(PIN) and ATP-Binding Cassette subfamily B (ABCB) proteins. In the presence of Bz-NAA, the signals of BODIPY-IAA2 in the inner part of the root were weak, while signals in epidermal cells was detected (Figure 7D, H, Supplemental movie7). These results suggest that the uptake of BODIPY-IAA2 into the lateral root cap and epidermis doesn’t require auxin transporter and that transport into the inner tissues is mediated by both auxin influx and efflux carriers. Both gravitropism and inhibitor assays suggest that BODIPY-IAA2 is transported by auxin transporters in the similar fashion to auxin. From these results, we concluded that our new fluorescent auxin with bright and stable property visualizes PAT mediated auxin transport, and thus could be used as a tool to study wide range of auxin-regulated biological processes.

**Figure 7.**
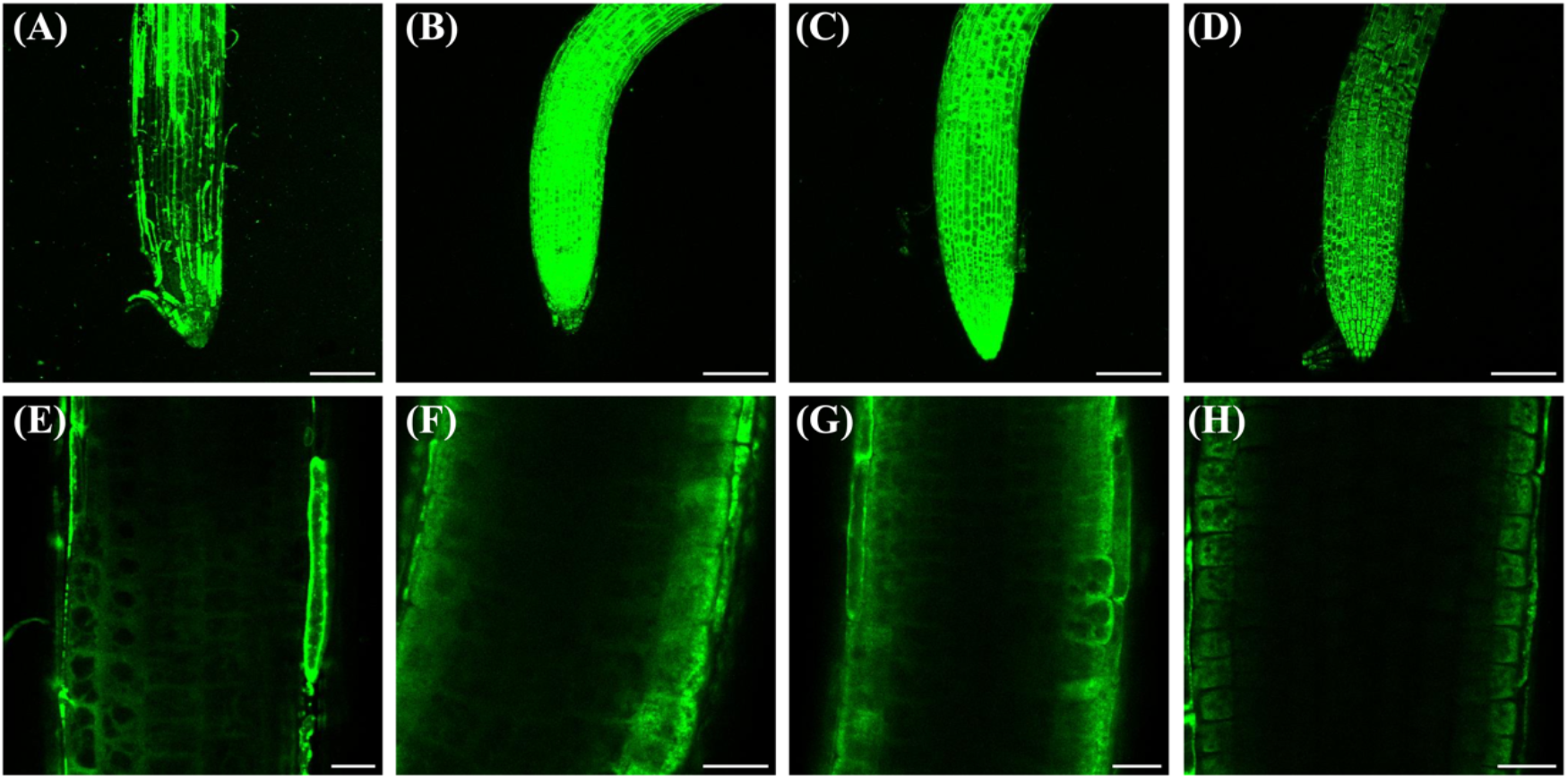
Distribution of BODIPY-IAA2 depends on auxin transporters. **(A-H)** Four-day-old *Arabidopsis* seedlings incubated with mock (A, E), 50 μM 1-NOA (B, F), 100 μM TIBA (C, G), and 50 μM Bz-NAA (D, H) for 1 hour and stained with 2 μM BODIPY-IAA2 for 30 min. Z-stack images were taken by confocal microscopy and maximum intensity projection images were shown (A-D). Scale bar: 100 μm (A-D), 20 μm (E-H).

## Discussion

Fluorescent auxin is a powerful tool for visualizing auxin dynamics in plants. As fluorescent auxin is a chemically modified auxin derivative, its physico-chemical property of the probe cannot be identical with that in auxin. Thus, careful consideration must be given to how well its staining pattern reflects the distribution of endogenous auxin. As direct visualization of endogenous auxin is currently impossible, one way is to compare staining patters of different fluorescent auxins. In this report, we developed BODIPY-IAA2 by replacing NBD part of NBD-IAA with BODIPY with optimization of the linker (Figure 1B, Hayashi et al. 2014). BODIPY is a fluorescent moiety suitable for live imaging, because it is very bright, has a short Stokes shift, is suitable for multicolor imaging, and is feasible for wavelength tuning. As with NBD-IAA, fluorescent moiety is attached to the position on IAA, modification of which results in loss of TIR receptor binding yet retaining activities to be transported by auxin transporters. This design was taken to the development of Bz-IAA, an auxin transport inhibitor, which also lost auxin signaling function while transported by auxin transporters (Tsuda et al. 2011). Our results showed that BODIPY-IAA2 inherit these attributes as well (Figure 2, 3, 6, 7). This is advantageous in visualizing auxin distribution without disturbing endogenous auxin function. On the other hand, the lack of auxin activity makes it difficult to correlate apparent localization pattern with auxin functions. However, this issue can be compensated by testing auxin transport inhibitors or loss-of-function mutants of auxin transporters. Indeed, our results suggest that BODIPY-IAA2 showed distribution patterns in *Arabidopsis* roots in an auxin influx and efflux carrier-dependent manner (Figure 7). Alternatively, having auxin signaling function, or anti-signaling function, in the probes is advantageous in correlating fluorescent pattern with phenotype, although it may also disturb auxin transporters by changing the level of their protein expressions (Sokołowska et al. 2014, Bieleszová et al. 2019, Bieleszová et al. 2024). Complementing these issues with different type of fluorescent auxin probes will be a challenge to visualize auxin dynamics in biological context.

Unlike proteins, visualizing sub-cellular localization of small-molecule phytohormones is difficult, and it indeed represents an outstanding question in auxin biology (Friml 2022). A major challenge is to visualize auxin distribution within a cell, which is difficult to achieve with current genetically encoded auxin sensors. For example, recently developed FRET auxin sensor with mid µM level sensitivity may not be insufficient to detect intra-cellular dynamics of auxin (Herud-Sikimić et al. 2021). Chemical modification of auxin is an alternative approach to the issue, and our results showed that BODIPY-IAA2 accumulated in the cytoplasm and ER. This subcellular distribution is also observed in other fluorescent auxins including NBD-IAA, NBD-NAA, and NBD-2.4-D (Hayashi et al. 2014, Pařízková et al. 2021). Thus, the results are robust as seen in both natural and synthetic auxins irrespective of the fluorescent moiety. The bright and stable property of BODIPY-IAA2 opened the door to analyze sub-cellular dynamics of auxin distribution in response to environmental stimulus. Comparative studies with other fluorescent auxins, including IAA-FITC retaining auxin activity, or the DNS-labelled IAA with anti-auxin activity, will clarify the role of sub-cellular localization in auxin functions.

## Materials and Methods

### Plant materials and culture conditions

*Arabidopsis thaliana* used in this study are the Col-0 ecotype. *Arabidopsis* seeds were sterilized and vernalized for 2-3 days at 4°C in the dark. Plants were germinated and cultured on 1ohormone auxin plays a crucial role in plant growth and response to envirμM MES (pH5.8), 0.8% (w/v) sucrose, 0.8% (w/v) agar) at 22°C under continuous white light conditions. The Cove-NIBB strain of *Physcomitrium patens* was used (Ashton and Cove, 1977). Plants were cultured on BCDAT (1 mM MgSO_4_, 1.84 mM KH_2_PO_4_, 10 mM KNO_3_, 45 μM FeSO_4_, 5 mM Ammonium Tartrate, 1 mM CaCl_2_, 0.22 μM CuSO_4_, 10 μM H_3_BO_3_, 0.23 μM CoCl_2_, 0.1 μM Na_2_MoO_4_, 0.19 μM ZnSO_4_, 2 μM MnCl_2_, 0.17 μM KI) agar medium at 25°C under continuous white light conditions. *P. viscosa* used in this study was harvested in Nagoya area in 2020. The seeds were plated on 1% agar plate and germinated at 16°C, moved to 1/2 MS media and incubated at 23°C for 1 week to observe distribution of BODIPY-IAA2.

### Spectrum Measurement

Ten μM of BODIPY-IAA2 was dissolved in DMSO and the spectrum of BODIPY-IAA2 were measured using SpectraMax iD5 (Molecular Devices). To obtain Emission spectra, excitation was performed at 350 nm and the fluorescence intensity between 440-640 nm was measured in 1 nm increments. For Excitation spectra, excitation was performed in 1 nm increments in the range of 400 nm to 530 nm, and the fluorescence intensity at 570 nm was measured. To compare fluorescence intensities, both 10 μM BODIPY-IAA2 and 100 μM NBD-IAA solution were excited at 440 nm and the fluorescence intensity between 480-700 nm was measured in 5 nm increments.

### Root elongation assay

Three-day-old *Arabidopsis* seedlings grown on vertical plates were transferred to 1/2 MS agar medium containing DMSO, 10 μM IAA, 10 μM BODIPY-IAA, 10 μM BODIPY-indole or 10 μM NBD-IAA. To analyze the interaction between IAA and fluorescent probes, 3-day-old *Arabidopsis* seedlings were transferred to 1/2 MS agar medium containing DMSO, 10 μM BODIPY-IAA, 10 μM BODIPY-indole or 10 μM NBD-IAA and 1 μM IAA. Then, plants were incubated on vertical plates for 24 hours at 22 °C under continuous white light conditions. Root elongation was calculated by measuring root length at the beginning of incubation and after 24 hours.

### GUS staining

*Arabidopsis* plants harboring DR5:GUS (Ulmasov et al. 1995) or BA3:GUS (Oono et al. 1998) were sown on 1/2MS medium containing 0.8% sucrose and 0.8% agar and grown at 22°C under continuous light for 6 days. The seedlings were put on 1/2MS agar medium containing 10 μM NBD-IAA, BODIPY-IAA or BODIPY-Indole and 5 μM IAA in f 60×15 mm petri dish (Greiner Bio One, Cat. No. 628 160) and petri dishes were put vertically and incubated at 22°C for 4 h. For examining the dose-dependent effects of BODIPY-IAA and BODIPY-Indole, seedlings were mounted onto 1/2MS medium containing 0, 1, 10 or 100 μM BODIPY-IAA (or BODIPY-Indole) and 5 μM IAA, and incubated at 22°C for 4 h. GUS staining was conducted according to Okamoto et al. 2013. Briefly, seedling samples were fixed with 90% ice-cold acetone overnight, washed with 50 mM phosphate buffer (pH 7.0) and incubated with 0.05% Tween 20 (Kanto Chemical Cat. No. 40350-02) in 50 mM phosphate buffer (pH 7.0) for 1 h. The samples were incubated with GUS staining buffer containing 2.5 mM potassium ferricyanide (Kanto Chemical Cat. No. 32337-30), 2.5 mM potassium ferrocyanide (Kanto Chemical Cat. No. 32338-30), 0.5 mM X-Gluc (Biosynth Carbosynth Cat. No. EB04274) in 50 mM phosphate buffer (pH 7.0) under vacuum for 20 min and incubated at 37°C for 2 h (DR5:GUS) or 6 h (BA3:GUS). The GUS reaction was stopped by adding 70% EtOH and the chlorophyll pigments were bleached. The bleached seedlings were mounted onto the slide glasses and taken images with 10x lens under the inverted microscope ECLIPSE TS2 (Nikon, Tokyo Japan).

### Yeast two-hybrid assay

Yeast strains harboring TIR1 and DII domain constructs were obtained from Dr. Naoyuki Uchida from Nagoya University. Yeast two-hybrid assays were performed as described previously (Uchida et al., 2018).

### Fluorescent probe staining

*Arabidopsis* seedlings were incubated with 1/2 MS liquid medium containing 1 μM BODIPY-IAA, 1 μM BODIPY-Indole or 1 μM NBD-IAA for 1 hour under dark conditions. Protonemal cells of *P. patens* were inoculated to BCD thin-layered agar medium on 3.5 cm glass bottom dishes and incubated for 5 days at 25°C under continuous white light conditions. Then, plants were incubated with 1/2 BCD liquid medium containing 1 μM BODIPY-IAA or 1 μM NBD-IAA for 1 hour under dark conditions. Seedlings of *P. viscosa* were incubated with 1 μM BODIPY-IAA solution for 1 hour under dark conditions.

### Root gravitropism assay

Five-day-old *Arabidopsis* seedlings grown on vertical plates were transferred to 1/2 MS agar medium containing 2 μM NPA, 10 μM NBD-IAA, 50 μM NBD-IAA, 10 μM BODIPY-IAA, 50 μM BODIPY-IAA, 10 μM BODIPY-Indole or 50 μM BODIPY-Indole. Each plate was rotated at angle of 90° and incubated in dark conditions. The bending angles of root tips were measured after 6 hours.

### Auxin transport inhibitor assay

Four-or 5-day-old *Arabidopsis* seedlings were incubated with 1/2 MS liquid medium containing Mock, 50 μM 1-NOA, 100 μM TIBA or 50 μM 1Bz-NAA for 1 h under dark conditions. Then, plants were stained with 2 μM BODIPY-IAA solution containing Mock, 50 μM 1-NOA, 100 μM TIBA or 50 μM 1Bz-NAA for 30 min under dark conditions. Stained samples were used for microscopic observation.

### Microscopy

Fluorescence images were obtained using confocal laser scanning microscopy (TCS SP8 FALCON, Leica) equipped with a white laser, a HyD detector and HC PL APO CS2 20×/0.75 objective lens. Plant cells stained with fluorescent probes were excited with 488 nm and their emission was acquired at 490-540 nm using a notch filter (488 nm).

## Supporting information

Supplemental movie 1. BODIPY-IAA staining in Arabidopsis root

Supplemental movie 2. BODIPY-Indole staining in Arabidopsis root

Supplemental movie 3. NBD-IAA staining in Arabidopsis root

Supplemental movie 4. BODIPY-IAA staining with Mock solution in Arabidopsis root

Supplemental movie 5. NBD-IAA staining with 1-NOA in Arabidopsis root

Supplemental movie 6. NBD-IAA staining with TIBA in Arabidopsis root

Supplemental movie 7. NBD-IAA staining with Bz-NAA in Arabidopsis root

## Acknowledgements and Funding

We thank Dr. Naoyuki Uchida in Nagoya University for providing Y2H strains and Miki Muranaka for technical assistance. This work was supported by the Japan Society for the Promotion of Science (JSPS) KAKENHI (21H04775 to Y.T.; JP22K21352, JP23H04739, JP22H04926 to Y.S; JP21K05068,JP24K21821 to M.N.), the Japan Science and Technology Agency (JST) Moonshot (JPMJMS2033-11) to Y.S., JST A-STEP(JPMJTR23UJ) to Y.S., The Naito Science & Engineering Foundation to T.A., and JST Core Research for Evolutionary Science and Technology (CREST) (JPMJCR1924). JSPS and NU are acknowledged for funding this research through the World Premier International Research Center Initiative (WPI) program.

---

**Supplemental Figure S1.**
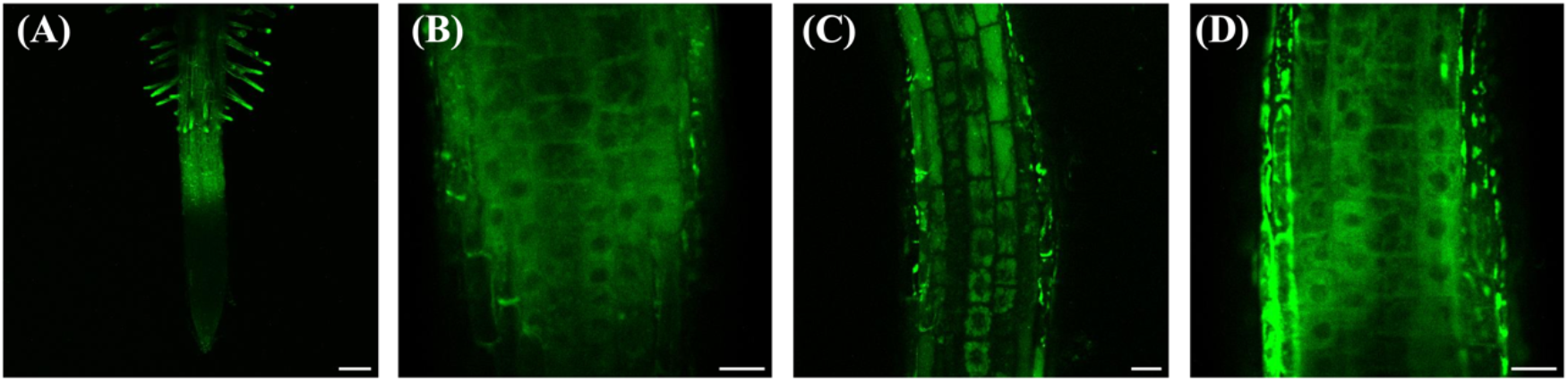
Staining patterns of fluorescent auxins. Staining patterns of NBD-IAA (A,B), TG-IAA(C), and BODIPY-IAA2(D) in root of *Arabidopsis*. Four or 5-day old seedlings of WT(Col-0) were incubated with 1 μM NBD-IAA (A,B), TG-IAA (C), and BODIPY-IAA2 (D). Scale bar: 100 μm (A), 20 μm (B-D)

**Supplemental Figure S2.**
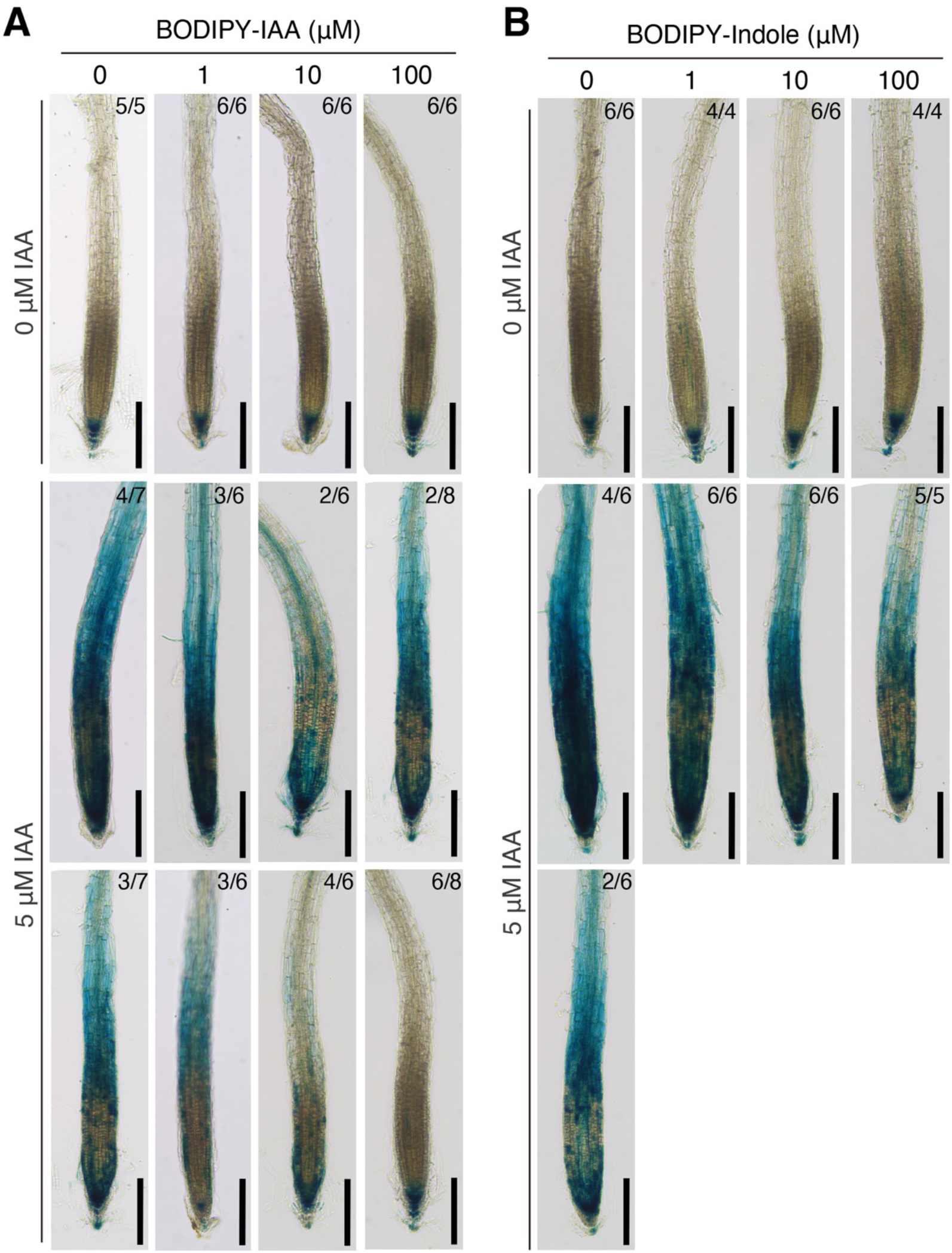
Dose-dependent effects of BODIPY-IAA and BODIPY-Indole on *Arabidopsis* DR5:GUS reporter line. Effects of BODIPY-IAA (A) and BODIPY-Indole (B) on auxin-responsive reporter gene expression. Six-day-old DR5:GUS seedlings incubated 0 or 5 μM IAA with 0, 1, 10 and 100 μM BODIPY-IAA or BODIPY-Indole for 4 h. Bar = 0.2 mm. Fractions on each image indicate the number of phenotypes observed (numerator) and the number of seedlings examined (denominator).

**Supplemental Figure S3.**
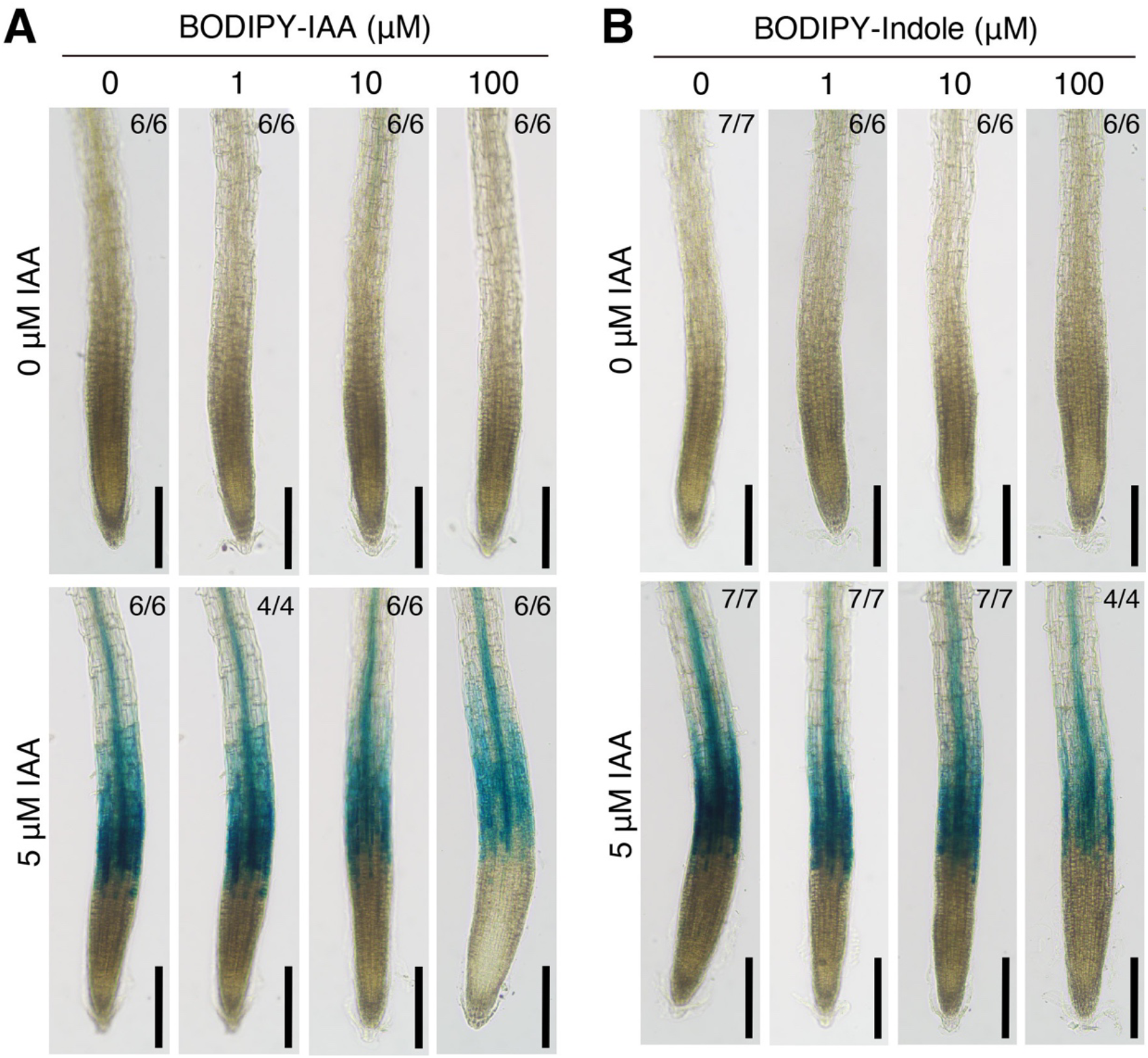
Dose-dependent effects of BODIPY-IAA and BODIPY-Indole on BA3:GUS reporter line. Effects of BODIPY-IAA (A) and BODIPY-Indole (B) on auxin-responsive reporter gene expression. Six-day-old BA3:GUS seedlings were incubated 0 or 5 μM IAA, together with 0, 1, 10 and 100 μM BODIPY-IAA or BODIPY-Indole for 4 h. Bar = 0.2 mm. Fractions on each image indicate the number of phenotypes observed (numerator) and the number of seedlings examined (denominator).

